# The viable but non-culturable (VBNC) state of *Campylobacter jejuni* lacks propagation ability in the mouse intestinal tract

**DOI:** 10.1101/2025.07.25.666711

**Authors:** Mizuki Tsuchida, Yurina Ohno, Akihiro Hirata, Yasuo Inoshima, Ayaka Okada

## Abstract

*Campylobacter jejuni* is a foodborne pathogen found worldwide. It can exist in a viable but non-culturable (VBNC) state under environmental stresses and cannot be cultured in this state. Whether *C. jejuni* retains its pathogenicity or causes food poisoning by resuscitation from the VBNC state remains unclear. This research aimed to evaluate whether *C. jejuni* in the VBNC state should be accounted for in prevention strategies against *Campylobacter* food poisoning. The VBNC state of *C. jejuni* strain NCTC11168 was induced at 4°C under aerobic conditions. A mouse model (C57BL/6, 3-week-old male) was prepared by administering a defined zinc-deficient diet and antibiotics in drinking water. Each mouse was administered *C. jejuni* with approximately 10^7^ colony-forming units of culturable cells, cells in the VBNC state, or Müller–Hinton broth as a control. In culturable cell-infected mice, an increase in the number of *C. jejuni* in stool was observed using colony counting and quantitative polymerase chain reaction. No culturable cells were present in VBNC cell-infected mice. Clinical signs, weight loss, or significant increase in lipocalin-2 and myeloperoxidase levels, as inflammatory biomarkers, were not observed in culturable cell- or VBNC cell-infected mice. Thus, culturable cells colonized the mouse intestinal tract, whereas cells in the VBNC state could not. In the VBNC state, *C. jejuni* may not pose a risk of food poisoning. Further investigations on the colonization and pathogenicity of *C. jejuni* in the VBNC state in the human intestinal tract are required.

## INTRODUCTION

*Campylobacter jejuni* is a gram-negative, microaerophilic bacterium causing gastroenteritis [13]. It is a foodborne pathogen distributed worldwide [17]. It causes watery or bloody diarrhea, fever, weight loss, headache, and nausea [26]. Most cases are associated with the consumption of *C. jejuni*-contaminated chicken meat and products [15].

A viable but non-culturable (VBNC) state has been identified in numerous bacteria, including *C. jejuni*. It is induced by various environmental stresses [34], such as low temperature [31] and distilled deionized water [7]. In the VBNC state, bacteria cannot grow in growth media; however, their metabolic abilities remain intact [4,10]. Previous studies have reported that some bacteria, such as *Salmonella* Typhi and *Vibrio parahaemolyticus,* in the VBNC state maintain their pathogenicity in the VBNC state [30,33]. Bacteria resuscitated from the VBNC state do not retain their pathogenicity in some cases; *V. vulnificus* in the VBNC state resuscitated in mice showed a lower ability to colonize the mouse gut, and was not pathogenic [1]. Because there are no reports on the pathogenicity of *C. jejuni* in the VBNC state, this should be investigated using an animal model. Although resuscitation from the VBNC state of *Campylobacter* has been reported in some studies, the resuscitation rate was extremely low [2]. Whether *C. jejuni* in the VBNC state retains its pathogenicity or causes food poisoning upon resuscitation from the VBNC state remains unclear.

Studies on the pathogenicity of *C. jejuni* have been largely limited owing to the lack of relevant and convenient animal models. Infection models, such as the deficient of single Ig interleukin (IL)-1 related receptor (Sigirr^−/−^) mice [27] and the anti-inflammatory cytokine IL-10 knockout (IL-10^−/−^) mice [20], have been reported. However, these models did not reflect acute bloody diarrhea [20]. To induce bloody diarrhea, a combination of IL-10 knockout and antibiotics for approximately two months is required [11]. Mice administered a defined zinc-deficient (dZD) food and antibiotics developed clinical signs in response to *C. jejuni* infection [9]. In this mouse model, *C. jejuni* infection induces both intestinal inflammation and systemic responses [9]. Genetic manipulations were not necessary, and the duration of antibiotic administration was reduced to two weeks. Therefore, we used a mouse model fed with a dZD diet and antibiotics to evaluate the pathogenicity and resuscitation of *C. jejuni* in the VBNC state.

## MATERIALS AND METHODS

### Animals

C57BL/6JJmsSlc (3-week-old male) mice were purchased from Japan SLC (Hamamatsu, Japan). The mice were maintained under controlled conditions at the Animal Experiment Facility of Gifu University. All animals were handled according to the regulations for animal experiments at Gifu University. All animal experiments were approved by the Animal Care and Use Committee of the Gifu University (approval number: AG-P-C-20230013).

### C. jejuni preparation

*C. jejuni* strain NCTC11168 was purchased from the American Type Culture Collection (ATCC, 700819, Manassas, VA, USA). The strain was stored in Brucella broth (211088, BD Biosciences, Billerica, MA, USA) containing 15% v/v glycerol (075-00616, Fujifilm Wako Pure Chemical, Osaka, Japan) at −80°C until use. The strain was cultured for two days on modified charcoal-cefoperazone-deoxycholate agar (mCCDA) (CM0739B, Oxoid, Hampshire, UK) with a CCDA- selective supplement (SR0155E, Oxoid) at 37°C under microaerobic conditions. To establish microaerobic conditions, an AnaeroPack-MicroAero agent (A-28; Mitsubishi Gas Chemical, Tokyo, Japan) was used in an anaerobic jar (A-110; Mitsubishi Gas Chemical). After two days of culturing, colonies were picked from mCCDA, suspended in 15 mL of Brucella broth, and cultured at 37°C for 40 hr under microaerobic conditions with shaking at 130 rpm in an incubator shaker (Innova 4300, New Brunswick Scientific, Enfield, CT, USA). After 40 hr of culture, 0.1 mL of liquid cultures from Brucella broth was resuspended in 15 mL Müller–Hinton (MH) broth (275730, BD Biosciences) and cultured at 37°C for 24 hr under microaerobic conditions with shaking at 130 rpm. The optical density of the liquid cultures at 600 nm was adjusted to 1.0, corresponding to a concentration of approximately 10^8^ colony forming units (CFU)/mL, using a spectrophotometer (GeneQuant 100, GE Healthcare, Chicago, IL, USA).

### Induction of the VBNC state

The VBNC state of *C. jejuni* was induced by culturing cells in MH broth at 4°C under aerobic conditions with shaking at 130 rpm for 26–28 days as described previously [32]. The induced VBNC cells were directly inoculated into mice without prior storage. The VBNC *C. jejuni* was cultured on Columbia blood agar base (CM0331, Oxoid) supplemented with 5% v/v defibrinated horse blood (Nippon Bio-Supp. Center, Tokyo, Japan) and maintained under microaerobic conditions at 37°C for 48 hr to assess the culturability. Cell viability was examined using the LIVE/DEAD *Bac*Light Bacterial Viability Kit (L13152, Invitrogen, Carlsbad, CA, USA), as reported previously [32].

### C. jejuni infection

The mice were provided with food and drinking water ad libitum. They were acclimated, fed a regular diet for three days, and then fed a dZD diet (0.00007% zinc, 17.33% protein) (CLEA Japan, Inc., Tokyo, Japan) for 14 days before infection. Prior to infection, mice were given an antibiotic mixture of gentamicin (35 mg/L, PHR1077, Sigma-Aldrich, MO, USA), vancomycin (45 mg/L, SBR00001, Sigma-Aldrich), metronidazole (215 mg/L, PHR1052, Supelco, MO, USA), and colistin (850 IU/mL, C4461, Sigma-Aldrich) in drinking water for three days as reported previously [9]. After three days of antibiotic administration, the mice were given sterilized water without antibiotics for one day before infection to eliminate the antibiotic effect on *C. jejuni* infection. Each mouse was administered approximately 1 × 10^7^ CFU of culturable or VBNC cells of *C. jejuni* in 0.1 mL of MH broth via oral gavage. Mock-infected mice were administered the MH broth alone. After administration, the mice were maintained on a dZD diet and sterilized water, and their body weight and clinical signs were measured at 1, 3, 5, and 7 days post-infection (dpi). Fecal samples were collected at 1, 3, 5, and 7 dpi. The mice were euthanized at 8 dpi. An overview of *C. jejuni* infections is presented in Fig. 1.

**Fig. 1.**
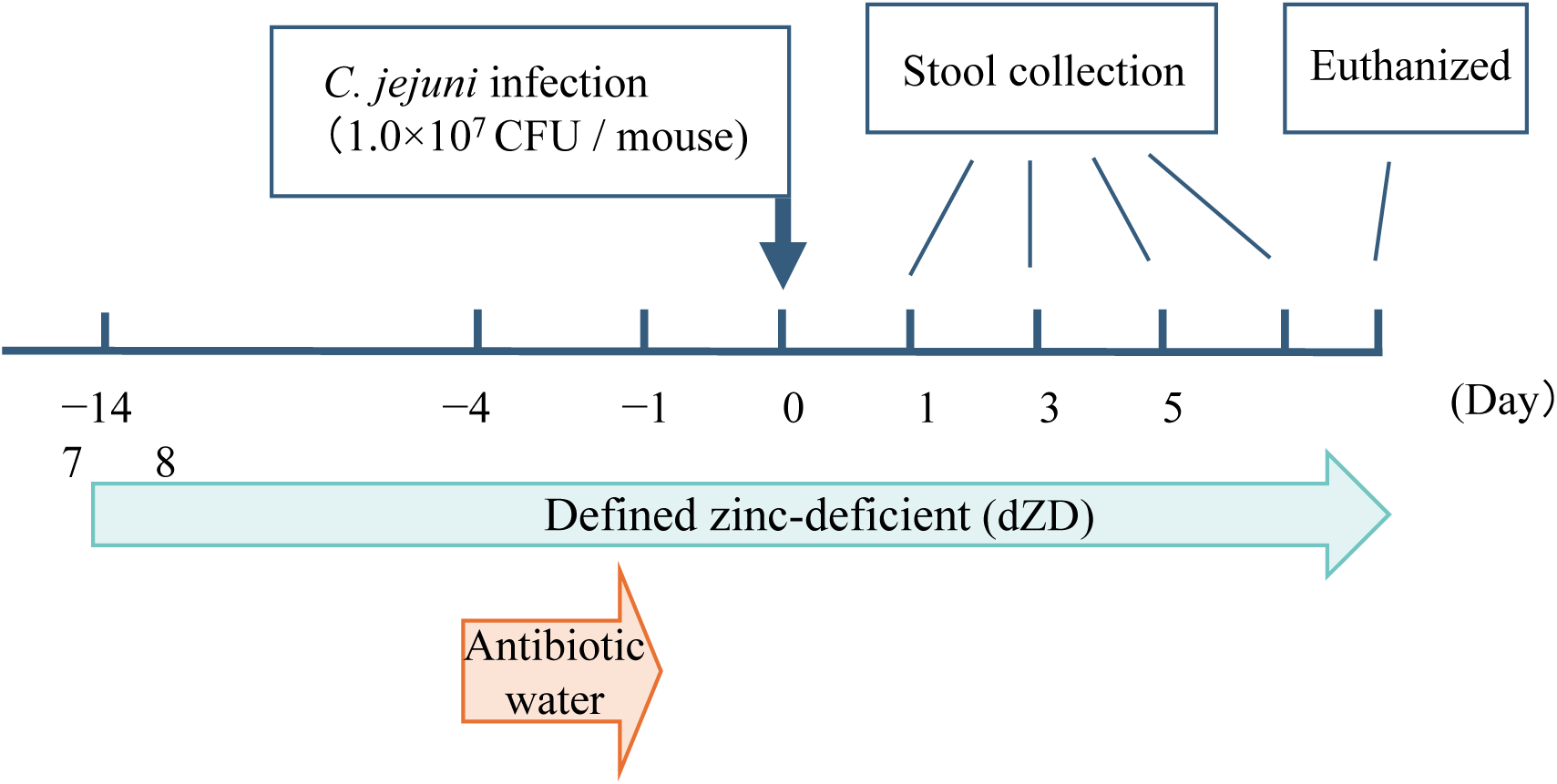
Schematic representation of the experimental design and sample collection. Mice were fed a dZD diet for 14 days before infection. They were administered an antibiotic mixture for 3 days and sterilized water without antibiotics for one day. The fecal samples were collected at 1, 3, 5, and 7 dpi. Mice were euthanized at 8 dpi. CFU, colony-forming units; dpi, days post-infection; dZD, zinc deficiency

### Sample collection

Fecal samples collected at 1, 3, 5, and 7 dpi were mixed with sterilized phosphate-buffered saline (PBS) and homogenized to obtain a 10% (w/v) fecal suspension. The fecal suspension was centrifuged at 800 × *g* for 10 min at 4°C, and the resulting supernatant was used for subsequent experiments. The supernatants were stored at −30°C until use.

### Culture-based enumeration

To enumerate culturable cells, 0.1 mL aliquots of 10-fold serial dilution of fecal supernatant were spread in duplicate on mCCDA with a CCDA-selective supplement and incubated at 37°C for two days under microaerobic conditions. A presumptive *Campylobacter* colony was selected from each sample and identified using polymerase chain reaction (PCR) with the C412F (5′-GGA TGA CAC TTT TCG GAG C-3′) and C1228R (5′-CAT TGT AGC ACG TGT GTC-3′) primers [18]. To detect damaged cells, approximately 0.05 mL of the fecal supernatant was mixed with 0.125 mL of Preston selective broth [(nutrient broth No. 2 (CM0067, Oxoid) base with 5% sterile lysed horse blood, *Campylobacter* growth supplement (SR00232E, Oxoid), and modified Preston *Campylobacter* selective supplement (6081, Condalab, Madrid, Spain)]. Samples were vortexed and incubated at 37°C for 24 hr under microaerobic conditions. Enrichment cultures were streaked on mCCDA with a CCDA-selective supplement and incubated at 37°C for 48 hr under microaerobic conditions.

### DNA extraction and quantitative PCR (qPCR) for C. jejuni quantification

DNA was extracted from 30 μL of fecal suspension using a DNeasy Blood & Tissue Kit (69506, Qiagen, Hilden, Germany) according to the manufacturer’s instructions. The elution volume was 30 μL. The extracted DNA was analyzed for the *C. jejuni/coli cadF* gene to determine the presence of *C. jejuni*, using primers for *cadF* (F, 5′-CTG CTA AAC CAT AGA AAT AAA ATT TCT CAC- 3′; R, 5′-CTT TGA AGG TAA TTT AGA TAT GGA TAA TCG-3′; Probe, 5′-FAM-CAT TTT GAC GAT TTT TGG CTT GA-BHQ1-3′) [9].

qPCR was performed using a Step One Plus thermal cycler (Applied Biosystems, Waltham, MA, USA) in a 96-well optical plate (Applied Biosystems) by interpolating the Ct values for each run with a standard curve of known amounts of *C. jejuni* DNA and transforming them into the number of organisms per milligram of fecal sample. The PCR mixture was made to a final volume of 20 μL containing 2 μL of the extracted DNA, using the GoTaq Probe qPCR Master Mix (Promega, Madison, WI, USA), according to the manufacturer’s instructions. Amplification cycle consisted of 5 min at 95°C, followed by 40 cycles of 15 s at 95°C and 60 s at 58°C [9].

### Quantification of lipocalin-2 (LCN-2) and myeloperoxidase (MPO) in fecal samples

The inflammatory status of mice was evaluated by measuring the levels of LCN-2 and MPO in fecal supernatants using DuoSet ELISA Mouse Lipocalin-2/NGAL (DY1857-05, R&D Systems, Minneapolis, MN, USA) and Mouse Myeloperoxidase DuoSet ELISA (DY3667, R&D Systems), respectively. Increased LCN-2 and MPO levels indicate epithelial damage [21] and neutrophilic inflammation [24], respectively. Fecal supernatants were centrifuged at 20,400 × *g* for 10 min at 4°C using an MX-301 high-speed refrigerated microcentrifuge (TOMY, Tokyo, Japan), and the resulting supernatant was used for ELISA. The supernatant was further diluted in PBS to a final concentration of 1:50 for LCN-2 or 1:10 for MPO and used for ELISA, according to the manufacturer’s instructions.

### Histopathological analysis

Ileum and colon segments were fixed in 4% paraformaldehyde, routinely processed, embedded in paraffin, cut into 5 μm sections, and stained with hematoxylin and eosin.

### Statistical analysis

Data analyses were performed using the EZR [16]. Differences among the three groups were evaluated using nonparametric statistics (Kruskal–Wallis). Differences were considered statistically significant at *p* < 0.05.

## RESULTS

### Culturability and viability of the administered C. jejuni

The VBNC cells were applied onto blood agar plates by spreading 100 μL bacterial suspension, which was adjusted to a concentration of approximately 1 × 10^8^ CFU/mL in culturable state before inducing the VBNC state. The number of observed colonies was less than 10 CFU/mL, and the bacterial viability was maintained at >90%. Bacteria administered to the mice were confirmed to be in the VBNC state.

### Clinical signs

The clinical signs of the mouse intestines were observed to characterize their response to *C. jejuni* every other day. Clinical signs, including inactivity, lack of responsiveness to stimulation, hunched posture, ruffled hair coat, soft feces, diarrhea, and weight loss were not observed in any mouse (Fig. 2). No difference in weight gain between mock-infected and *C. jejuni*-infected mice was noted (Fig. 2). Weight loss was observed 1–4 days before the challenge with the antibiotic mixture, which recovered after the intake of normal water. Therefore, the weight loss and clinical signs caused by *C. jejuni* were not observed in the present study.

**Fig. 2.**
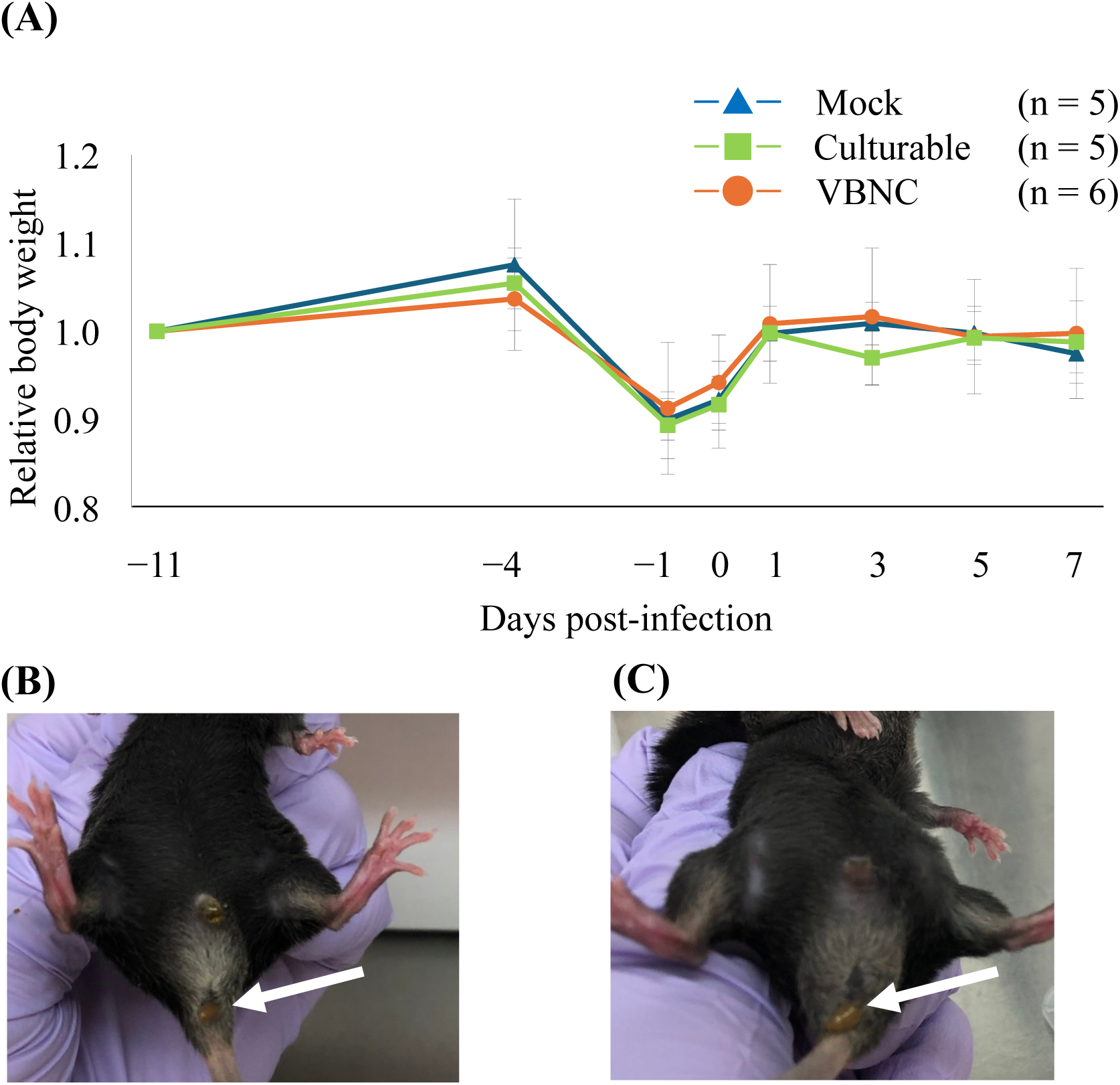
Changes in the body weight of mice and stools of mice as clinical signs. (A) The mean body weight for each group is expressed relative to the body weight of measured 11 days before infection (−11). The soft or normal stools were examined regardless of infection. (B) Mock-infected and (C) VBNC-infected group at 5 dpi. No significant difference was not observed. Arrows indicate stools. 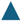, mock-infected (*n* = 5); 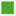, culturable cell (*n* = 5); 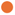, VBNC cell (*n* = 6). Error bars indicate standard errors. VBNC, viable but non-culturable dpi, days post-infection; VBNC, viable but non-culturable

### Quantification of culturable C. jejuni

Culturable *C. jejuni* in the fecal supernatant was counted using mCCDA with a CCDA-selective supplement (Fig. 3A). The colonies were identified as *Campylobacte*r spp. using PCR. *C. jejuni* was not detected at 1 dpi in either the culturable cell-infected or VBNC-infected groups. In the culturable cell-infected group, *C. jejuni* was detected in the stool of some mice at 3 and 5 dpi. Moreover, approximately 3.0 × 10^4^ CFU/10 mg feces of *C. jejuni* colonies were detected at 7 dpi in all mice in the culturable cell-infected group. However, no culturable *C. jejuni* were detected in the fecal samples in the VBNC cell-infected group. In addition, no resuscitation of VBNC cells was observed in the enrichment culture using the Preston selective broth (Table 1).

**Fig. 3.**
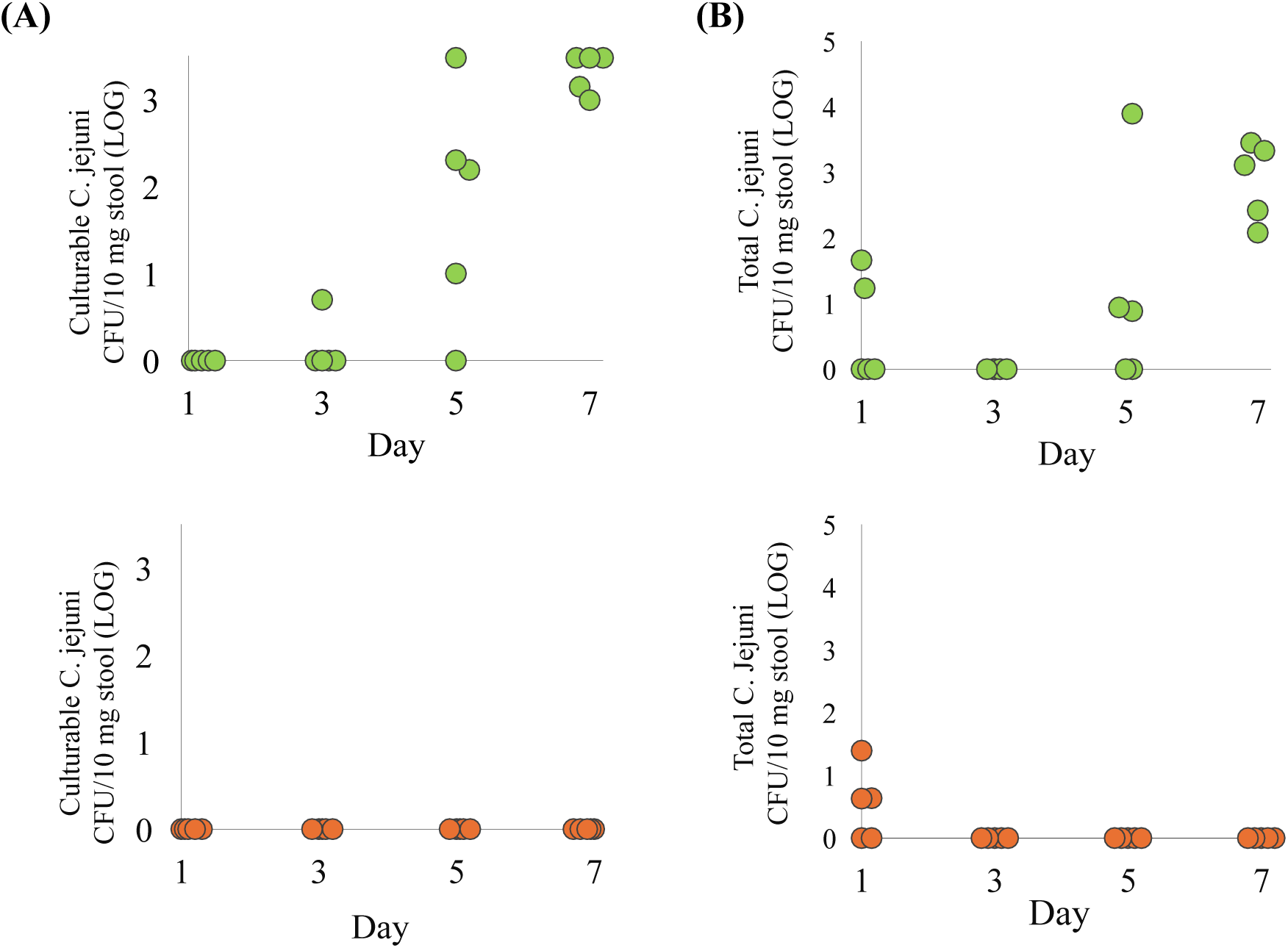
Culturable bacteria detected using mCCDA (A) and total bacteria detected using qPCR (B). The colonies on mCCDA were counted to quantify the bacteria in the feces and *Campylobacter jejuni* in the fecal supernatant was detected using qPCR to quantify total bacteria at 1, 3, 5, and 7 dpi. The upper panel shows the group infected with culturable cells, while the lower panel shows the group infected with VBNC cells. CFU, colony-forming unit; dpi, days post-infection; mCCDA, modified charcoal-cefoperazone-deoxycholate agar; qPCR, quantitive polymerase chain reaction; VBNC, viable but non-culturable

**Table 1.** Detection of culturable bacteria by Preston selective broth.

| Groups | Days post-infection |  |  |  |
| --- | --- | --- | --- | --- |
|  | 1 | 3 | 5 | 7 |
| Mock | 0/5 | 0/5 | 0/5 | 0/5 |
| Culturable | 1/6 | 0/6 | 4/6 | 5/6 |
| VBNC | 0/5 | 0/5 | 0/5 | 0/5 |

Results are expressed as the number of *C. jejuni* detected mice per number of total mice.
VBNC, viable but non-culturable

### Quantification of total C. jejuni using qPCR

The amount of *C. jejuni* including the non-culturable state, in the fecal samples was quantified using qPCR (Fig. 3B). *C. jejuni* was detected at 1 dpi in both groups, except in the mock-infected group. However, in mice infected with the VBNC cells, *C. jejuni* was not detected at 3 dpi. In mice infected with culturable cells, *C. jejuni* was not detected at 3 dpi but was detected again at 5 dpi in some mice. *C. jejuni* was detected in all the mice in the culturable cell-infected group at 7 dpi.

### Quantification of inflammatory biomarkers

The levels of inflammatory biomarkers, epithelial damage (LCN-2) and neutrophilic inflammation (MPO), in the fecal supernatant were measured at 1, 3, 5, and 7 dpi. No significant differences in LCN-2 and MPO levels were observed among all groups on any of the days tested (Fig. 4 and S1).

**Fig. 4.**
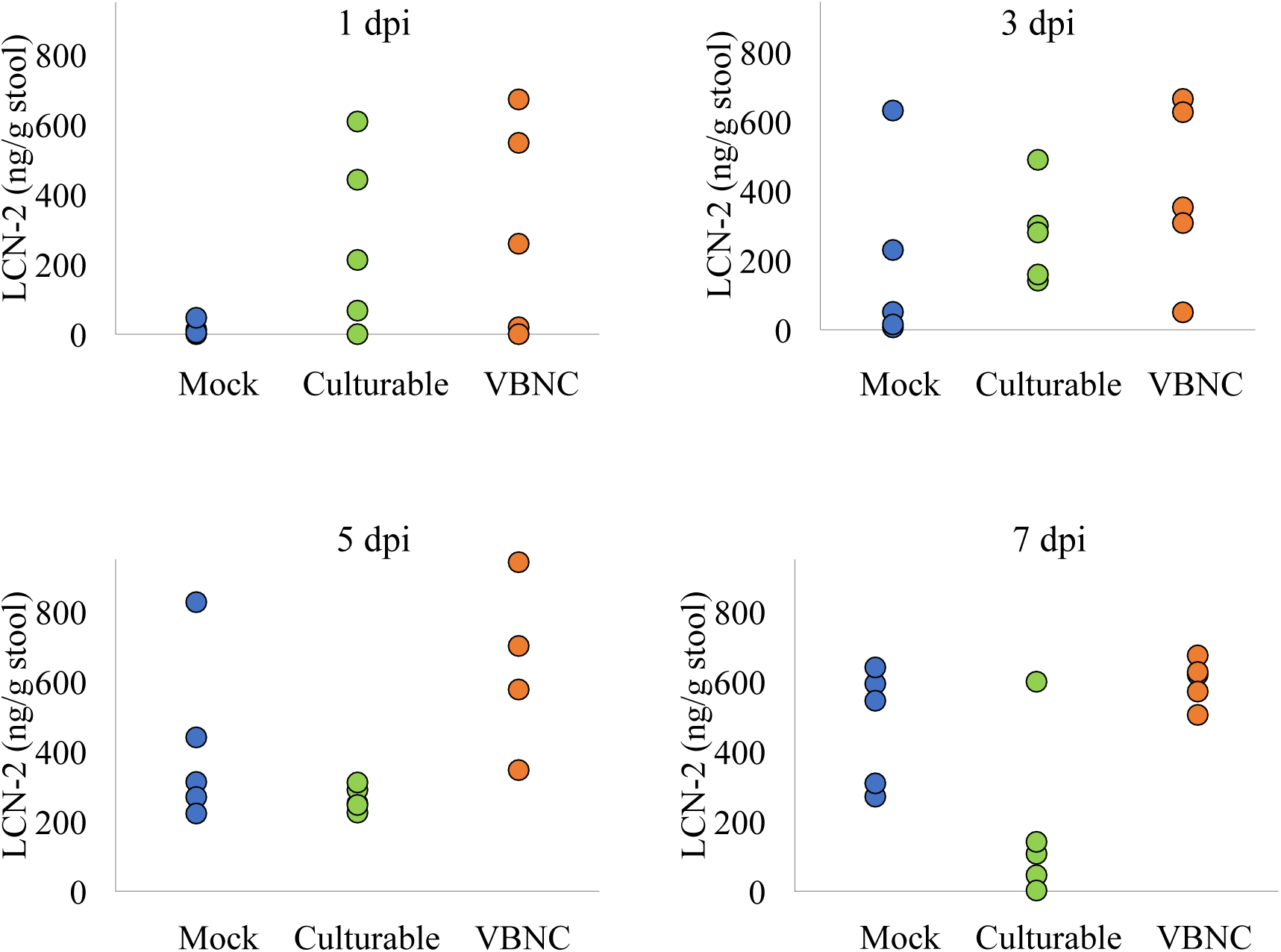
Quantification of LCN-2 levels in stool. LCN-2 levels in stool at 1, 3, 5, and 7 dpi. All groups consisted of *n* = 5, except for the 5 dpi group in which the VBNC cell-infected group had *n* = 4. No significant difference was observed. dpi, days post-infection; LCN-2, lipocalin-2; VBNC, viable but non-culturable

### Histopathological changes caused by Campylobacter infection

Cross-sectional slices were used to evaluate the effects of the infection of *C. jejuni* in the small intestine and the colon (Fig. 5). A small number of lymphocytes was observed in the lamina propria of the small intestine of some samples, regardless of whether they were infected. However, no substantial inflammation was observed in the small intestine and colon.

**Fig. 5.**
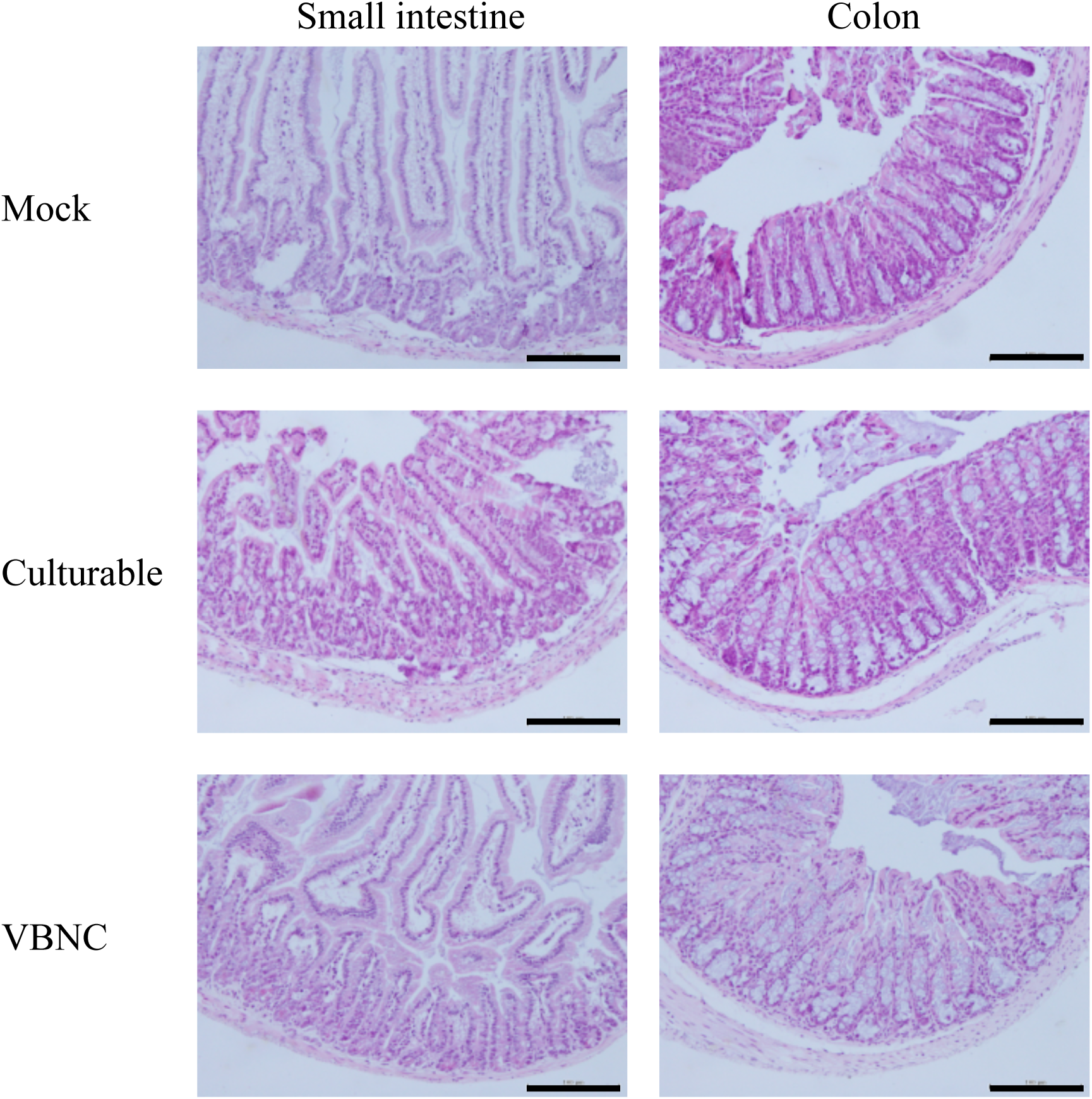
Histopathological observations. The small intestine and colon of mice in each group. No significant inflammation was noted. Scale bars indicate 100 μm. VBNC, viable but non-culturable.

## DISCUSSION

In this study, the effects of the infection of *C. jejuni* in the VBNC state on the mouse intestinal tract were evaluated. Host-specific commensal gut microbiota play an important role in the susceptibility or resistance of the vertebrate host to enteropathogenic infections in the intestinal tract [8]. High-dose *C. jejuni* infection does not exhibit pathogenicity in mice because of the physiological colonization resistance exerted by the murine gut microbiota composition [20]. Although *C. jejuni* colonizes the mouse intestinal tract, no clinical signs have been observed in normal mouse models [19]. Rodents are more resistant to inflammation than humans. Mice exhibit more than 10,000-times higher resistance to lipopolysaccharides and lipooligosaccharides, which are cell wall components of gram-negative bacteria [29]. Therefore, it is difficult to evaluate the inflammation caused by bacterial infection in a normal mouse model. In this study, a mouse model that showed colonization in the mouse intestinal tract and the same clinical signs observed in human campylobacteriosis was established by administering a dZD diet and antibiotics [9]. Although the same amount of culturable *C. jejuni* was administered as reported previously [9], clinical signs such as bloody stool and weight loss were not observed. LCN-2 and MPO levels in stool, which can increase without clinical signs, were measured to evaluate inflammation of the intestine [14]. LCN-2 reflects the extent of proinflammatory gene expression in the intestinal tract [6] and MPO is degranulated after migration to the gut mucosa during acute exacerbation [28]. These studies suggested that LCN-2 and MPO can be used to detect acute inflammation. However, no significant differences in LCN-2 and MPO levels were observed among the groups on any of the days tested in this study. The effects of antibiotics on the gut microbiota may not have been sufficient to cause the clinical signs in this experiment. The composition of the gut microbiota varies depending on the vendor [12], which may have weakened the effects of antibiotics in this experiment. Previous studies used the 81-176 strain [9]; however, we chose the NCTC11168 strain because inducing the VBNC state in the 81-176 strain requires significantly more time [32]. To address the possibility that strain differences may influence pathogenicity, we also conducted infection experiments using culturable 81-176. No clinical signs were observed in mice infected with 81-176 strain (data not shown). Moreover, even after administrating a tenfold higher dose of both 81-176 and NCTC11168 strains to the same mouse model, no clinical signs were observed (data not shown).

Although no clinical signs were observed, the colonization of *C. jejuni* in the mouse intestinal tract was detected. An increase in the number of *C. jejuni* in the fecal samples was detected in the culturable cell-infected group using colony counting and qPCR. The highest number of *C. jejuni* was detected using both methods at 7 dpi. However, an increase in the number of *C. jejuni* in the fecal samples was not detected in the VBNC cell-infected group by colony counting, and resuscitation was not detected. Using qPCR, *C. jejuni* in the fecal samples was only detected at 1 dpi and not detected at 3, 5, and 7 dpi. Therefore, *C. jejuni* in the VBNC state may not be able to colonize the mouse intestinal tract. To confirm the viability of *C. jejuni* in the VBNC state, we performed a CTC/DAPI staining to assess metabolic activity, in addition to the LIVE/DEAD staining, both of which indicated a high viability rate (Fig. S2). In our previous study, we also identified genes with significantly increased mRNA expression in bacteria induced into the VBNC state [22], further supporting the conclusion that the VBNC cells in this study were viable. Baffone et al. demonstrated that VBNC bacteria could be resuscitated in the mouse intestine [2]; however, the resuscitation rate was low, and no resuscitation occurred in long-term cultured samples. This suggests that the colonization may have resulted from a small number of residual culturable bacteria in the medium. Notably, we used C57BL/6JJmsSlc mice according to the previous report [9], whereas Baffone et al. used Balb/c mice, which may have contributed to the different results. Further studies are needed to clarify this isuue.

There are several possible reasons why the VBNC state *C. jejuni* failed to colonize. First, the ability of adhesins to bind fibronectin may be reduced. CadF plays an important role in adhesion to intestinal epithelial cells, and the expression levels of CadF decrease in the VBNC state [23]. Second, the ability of the flagellum may be reduced. The flagellum is considered to be a virulence factor [3]. In addition to its role in motility, the flagellum can transport bacterial effectors into the host cell and directly binds to the host cell receptor to trigger signaling involved in invasion [3]. In the scanning electron microscopy images, the loss of the flagellar structure of *C. jejuni* in the VBNC state was suggested [25]. Although the expression of *flaA* and *flaB*, which are associated with the flagellum, was maintained, downregulation of *flaA* and *flaB* was observed in the VBNC state [5]. Therefore, the downregulation of CadF and the flagellum may affect the colonization of the mouse intestinal tract in the VBNC state.

In this study, the clinical signs and inflammation of the intestinal tract caused by the infection of *C. jejuni* were not observed in either culturable or VBNC cell-infected mice. Therefore, the pathogenicity of *C. jejuni* in the VBNC state could not be evaluated. However, colonization of the mouse intestinal tract suggested that *C. jejuni* in the VBNC state lost its ability to colonize the mouse intestinal tract. Because in this study, we only evaluated colonization in the intestinal tract using one mouse strain, further investigations are needed to evaluate the resuscitation potential and pathogenicity of VBNC cells in both mouse and human intestinal tracts.

---

**Supplementary Figure 1.**
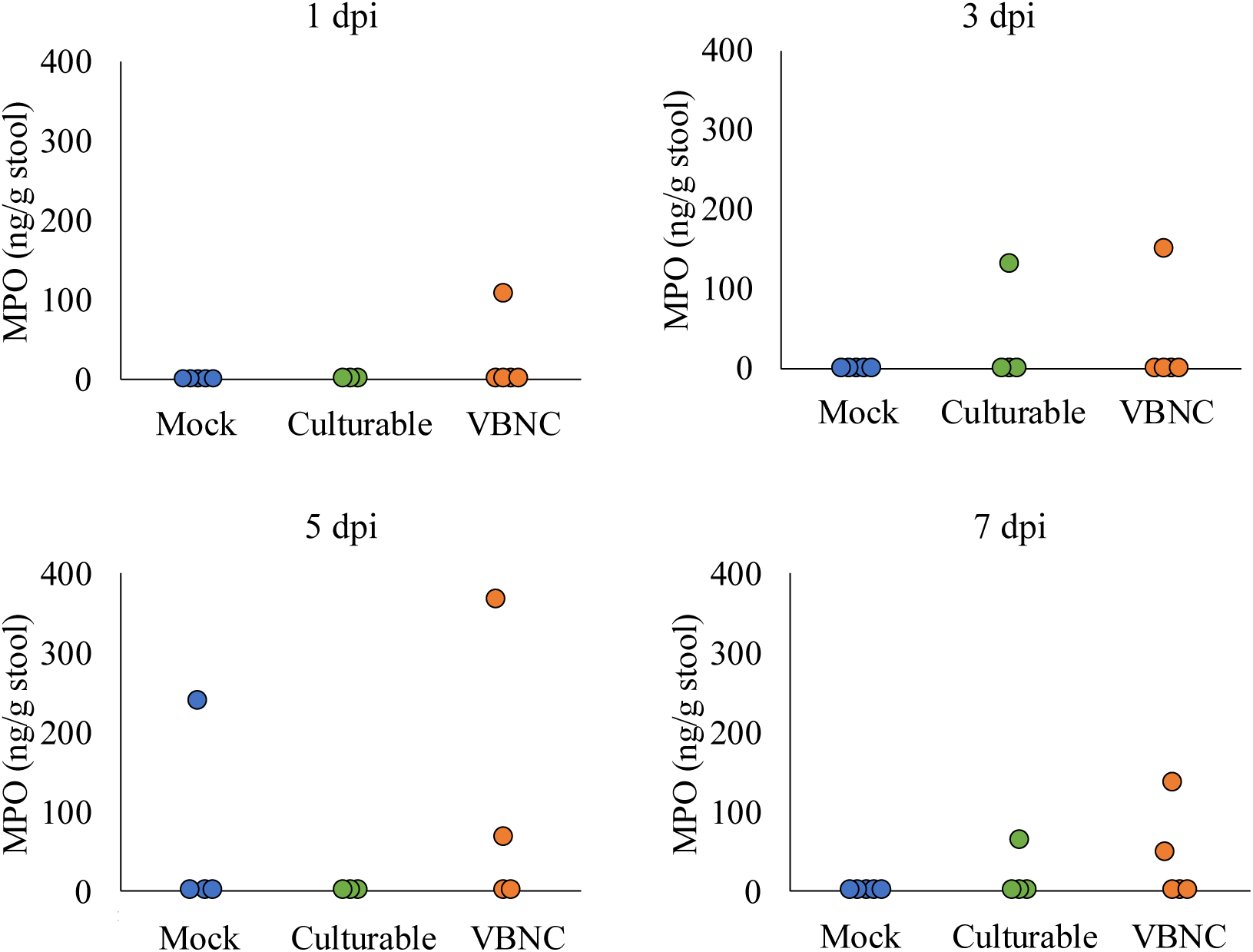
Quantification of MPO levels in stool. MPO levels in stool at 1, 3, 5, and 7 dpi. All groups consisted of *n* = 5, except for the 5 dpi group in which the VBNC cell-infected group had *n* = 4. dpi, days post-infection; MPO, myeloperoxidase; VBNC, viable but non-culturable

**Supplementary Figure 2.**
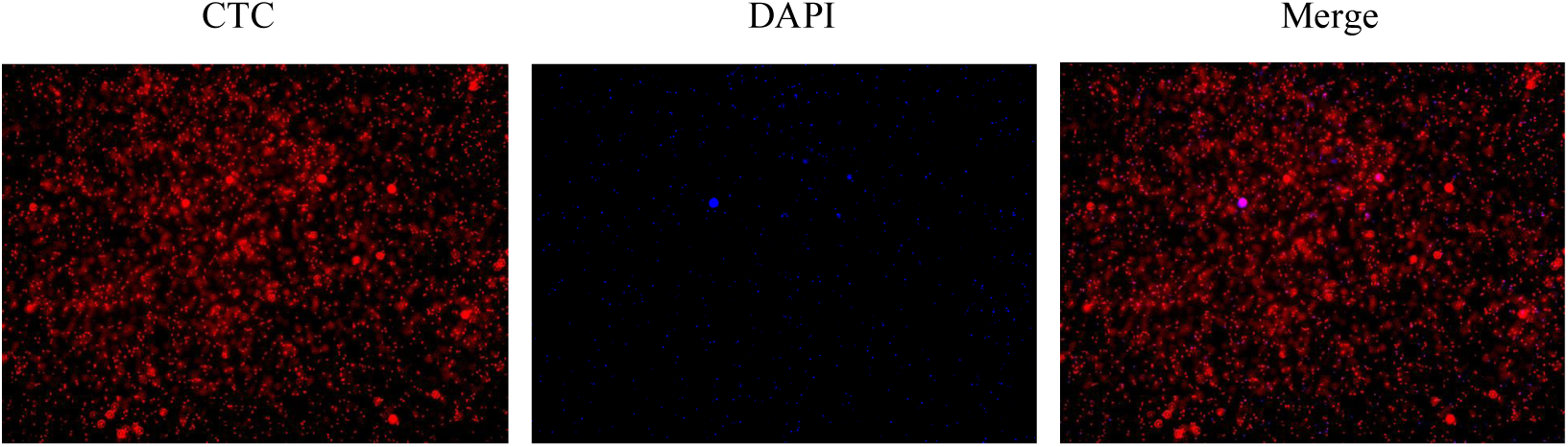
The images of CTC/DAPI staining of VBNC state of *C. jejuni* on day 28 induced using the same method used for mouse inoculation in this study. The red fluorescent signal is from CTCs, and the blue signal is from DAPI. CTC, 5-cyano-2,3-ditolyl-tetrazolium chloride; DAPI, 4’,6-diamino-2-phenylindole lactate; VBNC, viable but non-culturable

## REFERENCES

1. Baffone W, Citterio B, Vittoria E, Casaroli A, Campana R, Falzano L, Donelli G. 2003. Retention of virulence in viable but non-culturable halophilic *Vibrio* spp. Int J Food Microbiol 89: 31–39.

2. Baffone W, Casaroli A, Citterio B, Pierfelici L, Campana R, Vittoria E, Guaglianone E, Donelli G. 2006. *Campylobacter jejuni* loss of culturability in aqueous microcosms and ability to resuscitate in a mouse model. Int J Food Microbiol 107: 83–91.

3. Boehm M, Krause-Gruszczynska M, Rohde M, Tegtmeyer N, Takahashi S, Oyarzabal OA, Backert S. 2011. Major host factors involved in epithelial cell invasion of *Campylobacter jejuni*: role of fibronectin, integrin beta1, FAK, Tiam-1, and DOCK180 in activating Rho GTPase Rac1. Front Cell Infect Microbiol 1: 17.

4. Brenzinger S, van der Aart LT, van Wezel GP, Lacroix JM, Glatter T, Briegel A. 2019. Structural and proteomic changes in viable but non-culturable *Vibrio cholerae*. Front Microbiol 10: 793.

5. Chaisowwong W, Kusumoto A, Hashimoto M, Harada T, Maklon K, Kawamoto K. 2012. Physiological characterization of *Campylobacter jejuni* under cold stresses conditions: its potential for public threat. J Vet Med Sci 74: 43–50.

6. Chassaing B, Srinivasan G, Delgado MA, Young AN, Gewirtz AT, Vijay-Kumar M. 2012. Fecal lipocalin 2, a sensitive and broadly dynamic non-invasive biomarker for intestinal inflammation. PLoS One 7: e44328.

7. Chen H, Fu L, Luo L, Lu J, White WL, Hu Z. 2012. Induction and resuscitation of the viable but nonculturable state in a cyanobacteria-lysing bacterium isolated from cyanobacterial bloom. Microb Ecol 63: 64–73.

8. Fiebiger U, Bereswill S, Heimesaat MM. 2016. Dissecting the interplay between intestinal microbiota and host immunity in health and disease: Lessons learned from germfree and gnotobiotic animal models. Eur J Microbiol Immunol 6: 253–271.

9. Giallourou N, Medlock GL, Bolick DT, Medeiros PH, Ledwaba SE, Kolling GL, Tung K, Guerry P, Swann JR, Guerrant RL. 2018. A novel mouse model of *Campylobacter jejuni* enteropathy and diarrhea. PLoS Pathog 14: e1007083.

10. Hazeleger WC, Wouters JA, Rombouts FM, Abee T. 1998. Physiological activity of *Campylobacter jejuni* far below the minimal growth temperature. Appl Environ Microbiol 64: 3917–3922.

11. Heimesaat MM, Mousavi S, Bandick R, Bereswill S. 2022. *Campylobacter jejuni* infection induces acute enterocolitis in IL-10-/- mice pretreated with ampicillin plus sulbactam. Eur J Microbiol Immunol 12: 73–83.

12. Hufeldt MR, Nielsen DS, Vogensen FK, Midtvedt T, Hansen AK. 2010. Variation in the gut microbiota of laboratory mice is related to both genetic and environmental factors. Comp Med 60: 336–347.

13. Humphrey T, O’Brien S, Madsen M. 2007. Campylobacters as zoonotic pathogens: a food production perspective. Int J Food Microbiol 117: 237–257.

14. Jester BW, Zhao H, Gewe M, Adame T, Perruzza L, Bolick DT, Agosti J, Khuong N, Kuestner R, Gamble C, Cruickshank K, Ferrara J, Lim R, Paddock T, Brady C, Ertel S, Zhang M, Pollock A, Lee J, Xiong J, Tasch M, Saveria T, Doughty D, Marshall J, Carrieri D, Goetsch L, Dang J, Sanjaya N, Fletcher D, Martinez A, Kadis B, Sigmar K, Afreen E, Nguyen T, Randolph A, Taber A, Krzeszowski A, Robinett B, Volkin DB, Grassi F, Guerrant R, Takeuchi R, Finrow B, Behnke C, Roberts J. 2022. Development of spirulina for the manufacture and oral delivery of protein therapeutics. Nat Biotechnol 40: 956–964.

15. Kaakoush NO, Castano-Rodriguez N, Mitchell HM, Man SM. 2015. Global epidemiology of *Campylobacter* infection. Clin Microbiol Rev 28: 687–720.

16. Kanda Y. 2013. Investigation of the freely available easy-to-use software ‘EZR’ for medical statistics. Bone Marrow Transplant 48: 452–458.

17. Kirk MD, Pires SM, Black RE, Caipo M, Crump JA, Devleesschauwer B, Dopfer D, Fazil A, Fischer-Walker CL, Hald T, Hall AJ, Keddy KH, Lake RJ, Lanata CF, Torgerson PR, Havelaar AH, Angulo FJ. 2015. World Health Organization estimates of the global and regional disease burden of 22 foodborne bacterial, protozoal, and viral diseases, 2010: A data synthesis. PLoS Med 12: e1001921.

18. Linton D, Owen RJ, Stanley J. 1996. Rapid identification by PCR of the genus *Campylobacter* and of five *Campylobacter* species enteropathogenic for man and animals. Res Microbiol 147: 707–718.

19. Lone AG, Selinger LB, Uwiera RR, Xu Y, Inglis GD. 2013. *Campylobacter jejuni* colonization is associated with a dysbiosis in the cecal microbiota of mice in the absence of prominent inflammation. PLoS One 8: e75325.

20. Mansfield LS, Bell JA, Wilson DL, Murphy AJ, Elsheikha HM, Rathinam VA, Fierro BR, Linz JE, Young VB. 2007. C57BL/6 and congenic interleukin-10-deficient mice can serve as models of *Campylobacter jejuni* colonization and enteritis. Infect Immun 75: 1099–1115.

21. Moschen AR, Adolph TE, Gerner RR, Wieser V, Tilg H. 2017. Lipocalin-2: A master mediator of intestinal and metabolic inflammation. Trends Endocrinol Metab 28: 388–397.

22. Ohno Y, Rahman MM, Maruyama H, Inoshima Y, Okada A. 2024. Exploration of genes associated with induction of the viable but non-culturable state of *Campylobacter jejuni*. Arch Microbiol 206:260.

23. Patrone V, Campana R, Vallorani L, Dominici S, Federici S, Casadei L, Gioacchini AM, Stocchi V, Baffone W. 2013. CadF expression in *Campylobacter jejuni* strains incubated under low-temperature water microcosm conditions which induce the viable but non-culturable (VBNC) state. Antonie Van Leeuwenhoek 103: 979–988.

24. Prata MM, Havt A, Bolick DT, Pinkerton R, Lima A, Guerrant RL. 2016. Comparisons between myeloperoxidase, lactoferrin, calprotectin and lipocalin-2, as fecal biomarkers of intestinal inflammation in malnourished children. J Transl Sci 2: 134–139.

25. Santos LS, Rossi DA, Braz RF, Fonseca BB, Guidotti-Takeuchi M, Alves RN, Beletti ME, Almeida-Souza HO, Maia LP, Santos PS, de Souza JB, de Melo RT. 2023. Roles of viable but non-culturable state in the survival of *Campylobacter jejuni*. Front Cell Infect Microbiol 13: 1122450.

26. Skirrow MB. 1977. Campylobacter enteritis: a “new” disease. Br Med J 2: 9–11.

27. Stahl M, Ries J, Vermeulen J, Yang H, Sham HP, Crowley SM, Badayeva Y, Turvey SE, Gaynor EC, Li X, Vallance BA. 2014. A novel mouse model of *Campylobacter jejuni* gastroenteritis reveals key pro-inflammatory and tissue protective roles for Toll-like receptor signaling during infection. PLoS Pathog 10: e1004264.

28. Vermeire S, Van Assche G, Rutgeerts P. 2006. Laboratory markers in IBD: useful, magic, or unnecessary toys? Gut 55: 426–431.

29. Warren HS, Fitting C, Hoff E, Adib-Conquy M, Beasley-Topliffe L, Tesini B, Liang X, Valentine C, Hellman J, Hayden D, Cavaillon JM. 2010. Resilience to bacterial infection: difference between species could be due to proteins in serum. J Infect Dis 201: 223–232.

30. Wong HC, Shen CT, Chang CN, Lee YS, Oliver JD. 2004. Biochemical and virulence characterization of viable but nonculturable cells of *Vibrio parahaemolyticus*. J Food Prot 67: 2430–2435.

31. Wu B, Liang W, Kan B. 2016. Growth Phase, Oxygen, Temperature, and starvation affect the development of viable but non-culturable state of *Vibrio cholerae*. Front Microbiol 7: 404.

32. Yagi S, Okada A, Inoshima Y. 2022. Role of temperature, nutrition, oxygen, osmolality, and bacterial strain in inducing a viable but non-culturable state in *Campylobacter jejuni*. J Microbiol Methods 195: 106456.

33. Zeng B, Zhao G, Cao X, Yang Z, Wang C, Hou L. 2013. Formation and resuscitation of viable but nonculturable *Salmonella typhi*. Biomed Res Int 2013: 907170.

34. Zhang XH, Ahmad W, Zhu XY, Chen J, Austin B. 2021. Viable but nonculturable bacteria and their resuscitation: implications for cultivating uncultured marine microorganisms. Mar Life Sci Technol 3: 189–203.

